# QuantTB – A method to classify mixed *Mycobacterium tuberculosis* infections within whole genome sequencing data

**DOI:** 10.1101/676296

**Authors:** Christine Anyansi, Arlin Keo, Bruce Walker, Timothy J. Straub, Abigail L. Manson, Ashlee M. Earl, Thomas Abeel

**Author notes:** Corresponding author: Thomas Abeel.

## Abstract

**Background:** Mixed infections of *Mycobacterium tuberculosis*, and antibiotic heteroresistance, continue to complicate tuberculosis (TB) diagnosis and treatment. Detection of mixed infections has been limited to molecular genotyping techniques, which lack the sensitivity and resolution to accurately estimate the multiplicity of TB infections. In contrast, whole genome sequencing offers sensitive views of the genetic differences between strains of *M. tuberculosis* within a sample. Although metagenomic tools exist to classify strains in a metagenomic sample, most tools have been developed for more divergent species, and therefore cannot provide the sensitivity required to disentangle strains within closely related bacterial species such as *M. tuberculosis*.

Here we present QuantTB, a method to identify and quantify individual *M. tuberculosis* strains in whole genome sequencing data. QuantTB uses SNP markers to determine the combination of strains that best explain the allelic variation observed in a sample. QuantTB outputs a list of identified strains, their corresponding relative abundances, as well as a list of drugs for which resistance-conferring mutations (or heteroresistance) has been predicted within the sample.

**Results:** We show that QuantTB has a high degree of resolution, and is capable of differentiating communities differing by less than 25 SNPs and identifying strains down to 1× coverage. Using simulated data, we found QuantTB outperformed other metagenomic strain identification tools at detecting strains and quantifying strain multiplicity. In a real-world scenario, using a dataset of paired clinical isolates from a study of patients with either reinfections or relapses, we found that QuantTB could detect mixed infections and reinfections at rates concordant with a manually curated approach.

**Conclusion:** QuantTB can determine infection multiplicity, identify hetero-resistance patterns, enable differentiation between relapse and re-infection, and clarify transmission events across seemingly unrelated patients – even in low-coverage (1x) samples. QuantTB outperforms existing tools and promises to serve as a valuable resource for both clinicians and researchers working with clinical TB samples.

## Background

Tuberculosis (TB) - one of the oldest diseases in the world - continues to devastate the lives of millions per year. The World Health Organization’s *End TB Strategy* calls for a 95% reduction of TB deaths by 2035, a feat that will require more innovative and effective methods to treat, control and diagnose the disease (1).

For centuries it was assumed TB patients were infected with a single strain of *Mycobacterium tuberculosis,* the causative bacteria of TB. However, molecular genotyping methods have illuminated the phenomena of *mixed infections* - sometimes also referred to as superinfections or co-infections (2–6). Patients with mixed infections harbor multiple genetically distinct strains of TB at the same time. Previous research has suggested that mixed TB infections account for up to 30% of cases (4). However, the real incidence largely remains unknown (7), with estimates ranging from 19% for sputum samples up to 51% for combinations of pulmonary and extra-pulmonary samples (5). Mixed infections can complicate treatment and diagnosis through heteroresistance (presence of both drug susceptible and resistant patterns), which can cause false negatives in drug susceptibility tests and enable the spread of antibiotic resistance when left undetected (8–10). Therefore, accurate detection of strains within a mixed infection, as well as their distinct resistance patterns, is important for decreasing the worldwide TB burden and slowing the spread of drug resistance.

Various molecular typing methods have been used to gain clues as to whether a particular infection contains more than one *M. tuberculosis* strain. However, these approaches, which only examine a small portion of the genome, were not originally intended for the detection of mixed infections, and are limited in their sensitivity and ability to detect and quantify mixed infections. Restriction Fragment Length Polymorphism (RFLP) analysis relies on the positioning and copy number of the variable transposable insertion element IS6110 (11). Mycobacterial Interspersed Repetitive Unit-Variable Number Tandem Repeat (MIRU-VNTR) typing analyzes PCR amplified loci which vary in size and number of repeats (12). Finally, spoligotyping analyzes a series of 43 spacer oligonucleotides in the directed repeat region (12).

In contrast, whole genome sequencing (WGS) offers a more comprehensive view into the genetic composition of a sample, including distinct genetic information from individual strains. However, interpreting and analyzing such genomic data to identify and disentangle the composition of a mixed infection still remains a difficult task. To the best of our knowledge, no established methods exist to identify mixed infections for *M. tuberculosis* using WGS data. Some studies have classified a sample as mixed if the number of heterozygous positions (positions with evidence for more than one allele), exceeds a predefined arbitrary threshold (13, 14). These methods, which only consider mixes of two strains (bi-allelic variation), require sufficient coverage (>5x) for each allele and cannot be used to pinpoint actual strain identities. Other metagenomic tools exist to classify mixed populations of strains within a single species, such as Sigma, StrainEst, Strain Seeker, and Pathoscope (15–18); however these tools were developed and benchmarked using bacteria with greater intra-species diversity, such as *Escherichia coli,* where high numbers of variable sites and strain-specific structural variations can be exploited to delineate strains. These methods were not designed to be able to discriminate between strains of highly clonal species like *M. tuberculosis*, where there is near perfect syntenic gene conservation, and typically much less than 2000 genome wide SNPs between the most genetically distant isolates, resulting in an average sequence similarity over 99.97% between any two independent isolates.

We present QuantTB, a tool that is specifically designed to identify and quantify the abundance of closely related *M. tuberculosis* strains in WGS samples. QuantTB iteratively compares the SNPs from an uncharacterized TB sample with a database of TB SNP profiles from known reference strains, resulting in a low rate of false positives, while retaining sensitivity at coverages as little as 1x. Unlike other tools that were designed for use on species with higher levels of intra-species variation, QuantTB can accurately and precisely disentangle TB strains that differ by as few as 25 SNPs. QuantTB also informs the user of any drug resistant or hetero-resistant loci within the sample.

QuantTB is available on GitHub: https://github.com/AbeelLab/quanttb/

## Methods

### Construction of a SNP-based reference database

QuantTB uses a reference database of SNP sequences for strain classification which is constructed in four steps: 1) selecting a broad set of TB genomes, 2) selecting representative SNPs within these reference genomes 3) filtering genomes based on SNP similarity, 4) addressing reference genome bias.

#### 1. Acquiring genomes for the reference database

Although QuantTB can use either assemblies or raw sequencing reads for the construction of the reference database, assemblies are the preferred input. Assemblies represent aggregate, error-corrected versions of the corresponding read set and will yield superior results. We downloaded all available *M. tuberculosis* assemblies (5,867 complete and draft genomes as of July 23 2018) from NCBI (19, 20) using the taxonomic id: txid77643. We assigned lineages to each assembly based on lineage specific markers using a method described previously (21). We filtered out 217 assemblies that did not associate with any known *M. tuberculosis* lineage. We also removed 12 assemblies containing markers from more than one lineage. In total, 5,637 assemblies passed quality filtering. Supplementary Table 1 contains the NCBI accession codes and lineage prediction for all assemblies.

#### 2. Selecting representative SNPs

Selecting high quality SNPs for each genome present in the reference database is paramount to the success of our method. QuantTB can extract SNPs from two different sources: assemblies (FASTA files or SNP files outputted by MUMmer’s show-snps program (version 3) (22)) and read sets (FASTQ files or VCF files outputted by Pilon (version 1.22) (23)).

When extracting SNPs from assemblies, QuantTB aligns each assembly against the H37Rv reference genome (Genbank: CP003248.2) using MUMmer’s nucmer command with the minimum cluster length set to 100 (22) and other parameters set to the default values. All outputted SNPs are used, except for those marked as ambiguous by MUMmer. In the analysis presented here, we extracted SNPs from the 5,637 reference assemblies that passed quality filtering for our reference database.

Although not used for the analysis presented in this manuscript, QuantTB can also extract SNPs from read sets. QuantTB aligns each read set against the H37Rv (Genbank: CP003248.2) genome with BWA-MEM (Version: 0.7.17-r1188) (24) using default settings, then index-sorts with samtools (Version: 1.6, using htslib 1.6) (25). By default, QuantTB uses Pilon (version 1.22, fixes set to none) (23) to generate a pileup. Deletions, insertions, low coverage sites, low quality sites (Phred quality score less than 11), ambiguous sites (alternate allele frequencies less than 0.9), and reference calls are excluded.

For SNPs from both assemblies and read sets, we applied a number of additional filters. SNPs within a specified distance from one another (default 25bp) were removed from consideration, as these could be indicative of sequencing or alignment error. QuantTB also excludes all variants that are located in genes annotated as PE/PPE within the H37Rv reference, as these genes are known to be highly repetitive and prone to mapping errors, making it difficult to call variants using short-read data (26–28). The resulting SNP sequence for a genome is a dictionary of positions *(p)* that differ from the H37Rv genome mapped to their corresponding alleles, where *allele*(*p_*x*_*) → {*A, C, G, T*}. The complete collection of SNP sequences in the reference database is stored in a binary matrix, where rows are the genomes and columns are the locus/allele pair (Figure 1).

**Figure 1:**
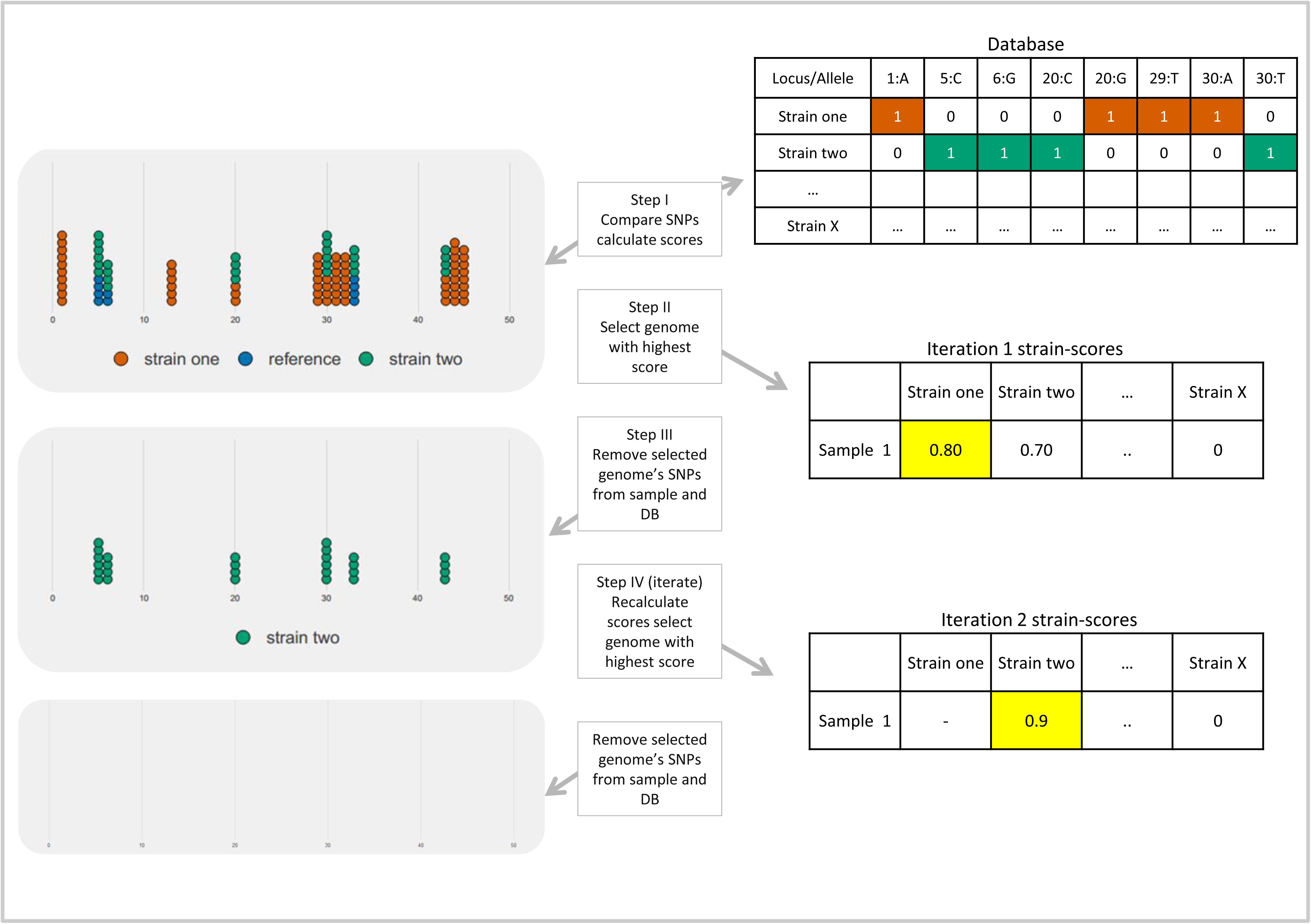
Iterative multiple strain identification process in QuantTB. First, SNPs from the sample are compared against SNP sequences in the reference database to calculate a strain presence score for every genome in the database. The sample is represented as a pileup, where every circle represents an allele copy. Red circles indicate alleles unique to strain A, green indicates alleles unique to strain B, and blue indicates reference strain (blue). The database (top right) is an example matrix representation of a reference genome database. Each column represents a single SNP (unique position and variant), and each row represents a genome in the reference database with this SNP present (1) or absent (0). Strain presence scores are calculated for every genome in the reference database. The genome with the highest strain presence score (*s_i_*) is selected, in this case strain A (red). The SNPs associated with strain A are removed from the database and the input sample, along with additional reference alleles. In each subsequent iteration the scores are recalculated, allowing for the identification of additional strains, and the process continues until there are no more SNPs or a threshold has been reached.

#### 3. Filtering genomes based on sequence similarity

The last step in constructing the reference database is to remove highly similar genomes. We calculated the pairwise SNP distances between each genome pair by summing the number of SNPs unique to each genome, i.e. by taking the union of variants minus the intersection of variants. If the SNP distance was below a specified threshold, the genome with the lowest number of SNPs was removed. This process was repeated until all genomes differed by the specified minimum SNP distance. We evaluated the performance of QuantTB by constructing reference databases with four different SNP distance thresholds: 10, 25, 50 and 100 SNPs. Table 1 shows the number of strains within each reference database.

**Table 1.**
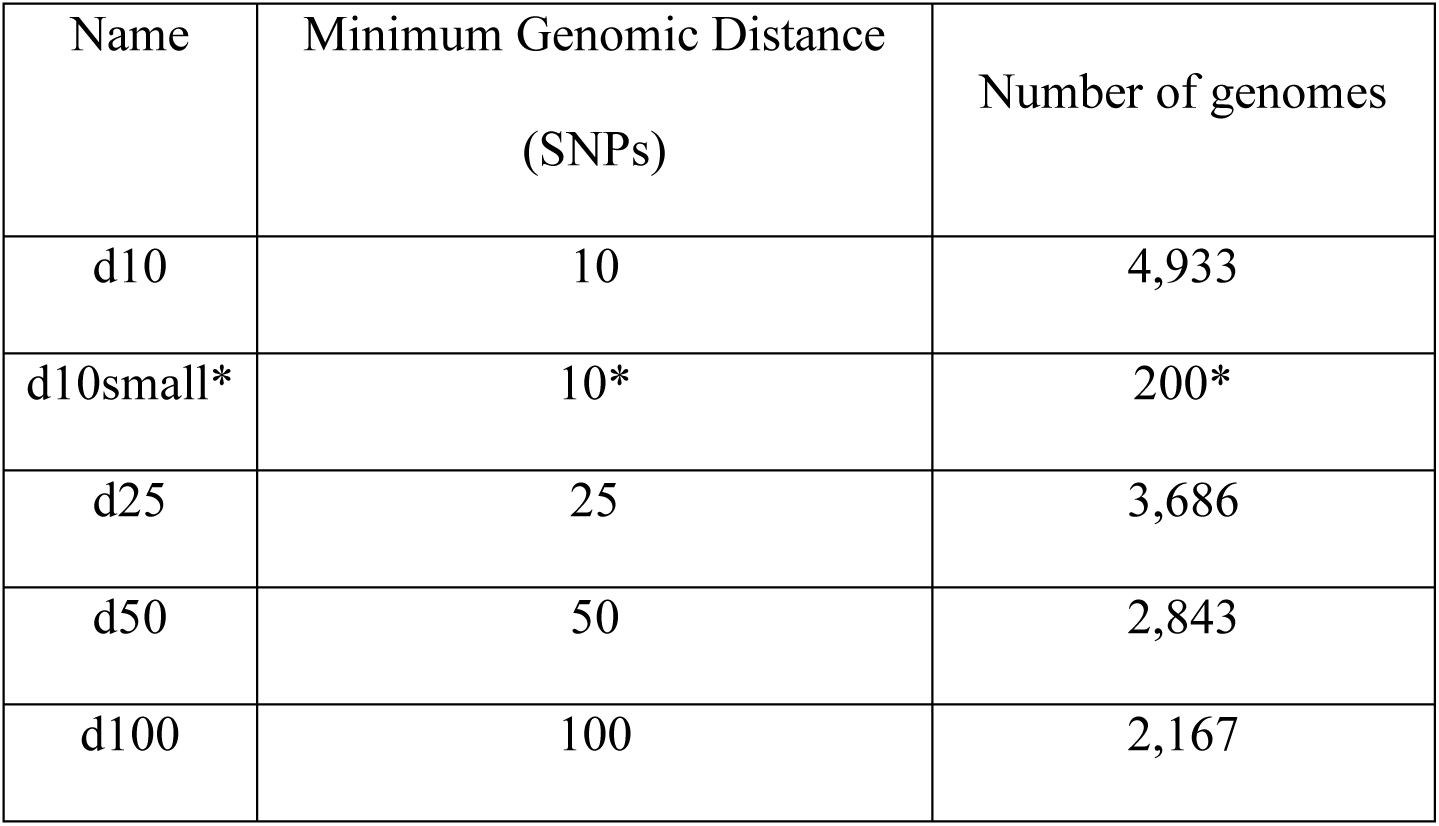
The number of genomes in each database after filtering by SNP distance. The distance was calculated by summing the number of unique SNPs between genomes. *In order to have a smaller database to benchmark against slower/more memory intensive tools, the number of genomes in d10small was restricted to be 200. The 200 genomes were randomly selected relative to the overall distribution of lineages, with a minimum requirement of five genomes for each lineage. D10 was selected as source set for the small benchmarking set to ensure the broadest possible strain and distance representation.

#### 4. Addressing reference genome bias

All SNPs were called using the reference genome, H37Rv, introducing a bias that strains highly similar to the reference genome become ‘invisible’ using this method, because they have a very low number of SNPs. To remedy this issue, a custom SNP-based representation of the H37Rv sequence was generated, based on the frequencies of SNPs across all other genomes in our reference database. If the same variant is observed in almost all the genomes in the reference database, we designate this as an H37Rv specific variant, i.e. a SNP within the H37Rv genome compared to every other genome. Therefore, QuantTB generates an “H37Rv SNP sequence” including positions where more than 75% of the genomes in the reference database have a common allele that differs from H37Rv. These locations are a fingerprint for H37Rv-like strains to identify them from the rest of the database.

### Using the SNP database to quantify strains present within a sample

QuantTB uses a SNP-based reference database to process short-read data in order to quantify the set of strain(s) present within a sample, such as short-read data from a clinical sample or isolate. Sample processing is done in two steps: 1) Extracting SNPs from a sample 2) Iterative classification of strains in the sample.

#### 1. Extracting SNPs from a sample

QuantTB can accept either a FASTQ file or a VCF file as an input sample for classification. Given a FASTQ file, reads are aligned against the H37Rv genome using BWA-MEM with default settings. A pileup is generated using Pilon with the default parameters and fixes set to none. Insertions, deletions, bases with low quality (Phred less than 11) and bases within PE/PPE regions are removed as in the construction in the reference database. All other bases with a frequency greater than 0.99 for the reference allele are removed. This process removes all of the reference H37Rv alleles, making it impossible to detect the H37Rv strain in a sample, as they have been filtered out. Therefore, to ensure H37Rv can be detected, we ensure all bases within the “H37Rv SNP sequence” are included (see construction of reference database above), given they pass the initial quality filter. The end result is a dictionary containing the extracted allele coverages and frequencies for every SNP position identified in the database. Note that QuantTB does not filter based on coverage; this allows for the detection of low abundance strains within a sample.

#### 2. Iterative classification of strains in the sample

Specific TB strains within the reference database are identified as present within a sample by iteratively querying against the SNP-based reference database as follows (Figure 1):

I. Compute a “strain presence score” (*s_i_*) for every genome *(i)* in the database (see below for computation of score).
II. Choose the genome with the highest strain presence score, *s_i_*.
III. Remove the chosen genome’s SNPs from the database and sample.
IV. Repeat steps 1-3 until no more SNPs remain, the strain presence score is below the threshold, or the maximum number of iterations have been reached.

##### Computation of strain presence score

During each iteration, a strain presence score *(s_i_*) is calculated for every genome in the database (*D).* The strain presence score is an average of two statistics, *O*_*i*_ and *A*_*i*_, and represents the overall presence of a strain within the sample. *O*_*i*_ and *A*_*i*_ are described below.

***O*_*i*_** represents the fraction of SNPs from a particular reference genome, *i*, that was observed in the sample. The higher *O_i_*, the more likely the set of SNPs observed in the sample originated from genome *i*.

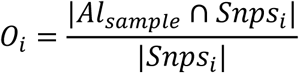

*Al*_*sample*_ is the set of alleles observed above a coverage threshold *t_a_*. Applying a coverage threshold diminishes the effect of random errors in the sample, while retaining sensitivity for true variation. This threshold *t_a_*, is dynamic and determined by the average coverage of the sample, *C_sample_*, and the average coverage of the genome identified in the previous iteration, *C*_*G*_*k* − 1__.

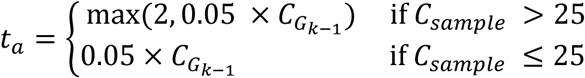

If the sample has an average coverage greater than 25, a minimum coverage threshold of 2 is set for all iterations, whereas for samples with an average coverage less than 25, there is no minimum, so that strains at low coverage can still be detected. For each iteration *k*, the threshold is set as 5% of the average coverage of the strain identified in the previous iteration. This is initialized at *k=0* as 5% of the sample coverage (*C_sample_*). Applying a coverage threshold diminishes the effect of random errors in the sample, while retaining sensitivity for true variation. Notice that this threshold likely goes down in every iteration as the coverage of the previously detected strain is used with a minimum of 2.

***A*_*i*_** represents the frequency with which a particular genome’s SNPs accounts for all the allelic variants present in the sample. The previous statistic, *O*_*i*_, represents how many SNPs of a particular genome been observed with sufficiently high coverage. However, when a sample has low coverage, the probability of observing the complete set of a genome’s SNPs is low. To account for strains present at low coverages, QuantTB also calculates, *A*_*i*_.

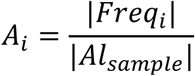

Where *Freq*_*i*_ represents the vector of frequencies for each allele of genome *i* within the sample:

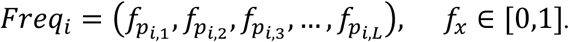

##### Choose the genome with the highest strain presence score

At the end of each iteration, the strain presence score *(s_i_,)*, is calculated as an average between *O_i_*and *A_i_*, and the genome with the highest *s*_*i*_, is selected as being present in the sample.

##### Remove the chosen genome’s SNPs from the database and sample

Before the next iteration begins, SNPs corresponding to the chosen genome are 1) removed from each SNP sequence in the database and 2) removed from the sample. In addition, any H37Rv alleles present in the sample at positions outside of the identified genomes’ SNP sequences are also removed. This is because those alleles can be accounted for already by the presence of the identified genome.

Because it is unlikely that the true strain present in the sample shares the exact collection of SNPs with its highest scoring match in the database, additional SNPs from the sample could match erroneously across multiple other genomes in the database with enough coverage to be marked as ‘observed’. As the coverage increases, the probability that an additional genome is spuriously detected also increases, due to the number of these uninformative SNPs that do not match perfectly with the originally selected genome. QuantTB implements a check to safeguard against this. To account for spuriously detected genomes due to higher coverages (greater than 25), we only allow strains to be detected in a sample when their prevalence accounts for at least 1% of the sample coverage. Therefore, SNPs from a particular strain are only removed from the sample when the change of coverage at each iteration would be at least 1%, otherwise the strain is ruled out for detection.

##### Iteration

The QuantTB algorithm iterates until one of the following stop criteria are reached: 1) the number of SNPs in the sample is below 15, 2) no genomes have more than 15 overlapping SNPs with the sample, or 3) the score threshold has been reached (the default is 0.15 but this can be adjusted by the user). At the end of the iterations, relative abundance is calculated by taking the average coverage of unique SNPs for each genome in the sample.

### Prediction of antibiotic resistance status of detected strains

In order to identify presence or absence of a resistance phenotype in the sample, QuantTB uses a curated set of SNPs conferring antibiotic resistance to seven TB drugs (21) (Supplementary Table 2). If resistance conferring allele(s) are present at a frequency of more than 90%, the sample is considered fully resistant for that drug. Heteroresistance, where there is evidence of both a resistant and a susceptible phenotype in a sample, can occur due to mixed infections or through in-host microevolution. If a resistance conferring allele(s) is present at a frequency between 10-90%, then the sample is considered heteroresistant for that drug. QuantTB outputs the results of the resistance testing in a separate file, if the appropriate command-line flag is set.

### Benchmarking using synthetic read sets

We constructed test datasets to benchmark QuantTB and compare its performance to two other strain level identification methods, StrainSeeker (16) and Sigma (15). Another tool, StrainEst (29) is also capable of performing single strain classification; however, a downloadable script is not provided to construct a database for *M. tuberculosis* genomes compatible with their algorithm, so we were unable to include it in our benchmark.

Synthetic mixed samples of two and four strains were used to perform benchmarking. In order to benchmark overall performance across different coverage levels, as well as across databases with different levels of strain similarity, we constructed mixes of four strains, where all four strains were present at equal relative abundance. In order to further benchmark the ability of QuantTB to assess samples containing strains with different relative abundances, we generated synthetic mixes of two strains sampled at different relative abundances.

To generate the four strain mixtures we randomly selected 200 combinations of four assemblies from each of the four reference databases generated with different SNP-distances using publicly available *M. tuberculosis* assemblies. In total, we selected 800 different combinations of four strains. For each reference database, we ensured that all 7 main lineages were represented across the selected sets of assemblies. Then, for each selected assembly, we synthesized paired end reads using ART (Version 2.5.8) (30) with default settings for the Illumina HiSeq 2500 platform, at a read length of 101 bp and a final coverage of 100×. Each read set was down sampled to 0.1×, 1×, 10×, and 20× coverage, then merged into mixes of four. This corresponds to 800 mixed sets of four different coverage levels, or 3200 synthetic mixes of strains.

To generate synthetic two-strain mixtures of strains at different relative abundances, we randomly selected 100 pairs of assemblies from each of the d50 and d100 reference databases. Paired end reads were simulated for each assembly, then the read sets were merged in mixes at 1×/9× coverage and 3×/7× coverage. This corresponds to 200 mixed sets at two different coverage levels, resulting in 400 synthetic mixes of varying relative abundance.

In addition, we generated synthetic four-strain mixtures for a smaller dataset, able to run in shorter compute time. StrainSeeker and Sigma are not capable of processing large sized reference sets (>2000 genomes) and required >3 days of compute time per sample or >7 days for reference database construction of 2,000 genomes. Therefore, to compare the performance of QuantTB against that of StrainSeeker and Sigma within a reasonable time frame, we created a smaller reference database, d10small. Using the reference genomes from the d10 database (see Methods), we randomly selected 200 genomes such that each TB lineage was represented in proportion to its relative incidence in the overall dataset, with a minimum requirement of five representatives for each lineage. Synthetic sample sets were then created based on the small reference set, using 200 randomly selected sets of 4 genomes. These sets were synthesized using the same method as for the previous databases, with the only exception being that we only created samples where the strains are present at either 1× and 10× coverage.

### Benchmark evaluation using synthetic sets

In order to test the performance of each method, we calculated the *Recall*, *Precision*, and the *F-measure* for every test category. True positive (*TP*) refers to the number of correctly identified strains. False positive (*FP*) refers to the number of identified strains that were not present in the sample. False negative (*FN*) refers to the number of strains present in the sample that were not identified.

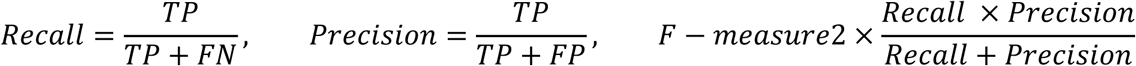

### Evaluation using real genomic data

We demonstrated the utility of QuantTB with real data samples from a study investigating reinfection and relapse using WGS (13). Sequencing reads from 50 pairs of isolates were downloaded from the SRA (31). SRA files were extracted using fastqdump (Version 2.9.0) (31) from the SRA toolkit, using the “split-3”, “skip-technical”, and “clip” flags to split left and right reads into separate files, remove technical reads, and clip off poor-quality ends of reads, respectively.

To construct a phylogenetic tree from these samples, SNPs were extracted and filtered as described above. FastTree (32) was used to generate a tree from the concatenated SNPs.

## Results

### Comprehensive TB reference database captures the breadth of the *Mycobacterium tuberculosis* species

QuantTB requires a reference database of known *M. tuberculosis* genomes for classification, where every genome is represented by a set of SNPs (see right panel in Figure 1). To generate a TB reference database, we downloaded all 5,867 complete and draft *M. tuberculosis* assemblies available at NCBI on July 23 2018 (see Methods). To remove non-TB genomes and possible chimeric assemblies, we assigned lineages to each assembly using a method described previously (21). After additional filtering on assembly quality and removing non-TB genomes, or genomes with more than one lineage classification, 5,637 assemblies remained.

Our database contained eight major lineages of TB at frequencies reflecting the overall abundances of sequences for each lineage in NCBI (Figure 2A). Lineage 4 strains encompass the vast majority of *M. tuberculosis* assemblies currently available at NCBI (3,455 strains), while lineage 7 and lineage 5 are the least abundant with 6 strains for each (Figure 2A). The genetic diversity within lineages (Figure 2b) was in agreement with previous studies (33): (i) lineage 1 had the greatest intra-lineage genetic diversity (median of 871 SNPs pairwise distance) and (ii) lineage 2, the second most frequently occurring lineage, had the lowest diversity, (median of 240 SNPs pairwise distance). The six strains that comprise lineage 7 had a wide range of genetic diversity, suggesting the need for increased sequencing of less well-characterized lineages, which would improve the resolution of classification within these less abundant lineages.

**Figure 2:**
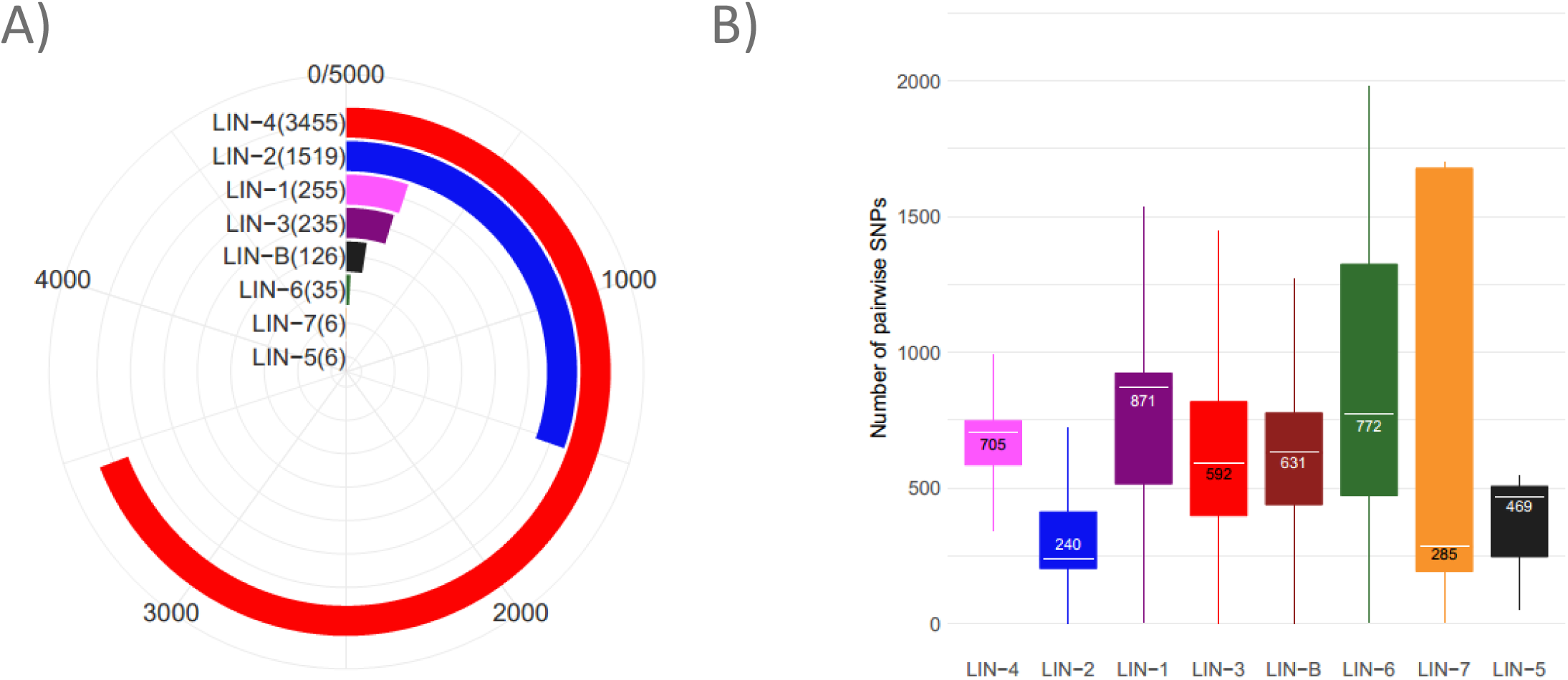
Panel A) Number of representatives from each lineage amongst all 5,637 *M. tuberculosis* assemblies in our reference database. Panel B) Intra-lineage pairwise distance for each lineage as measured by the number of unique SNPs between a pair. The number in the box plot is the median distance of all pairs of samples from that lineage.

To benchmark QuantTB’s performance across databases with varying intra-database genetic distances, we constructed a set of databases with differing minimum differences between strains. After extracting high quality SNPs from each genome in our database, we calculated pairwise SNP distances between genomes (see Methods) and filtered highly similar genomes using four different SNP distance thresholds (10, 25, 50, and 100 SNPs) into five databases (Table 1, Methods). Each database contained a representative distribution of strains from each lineage (Supplementary Table 3), as well as representative genetic diversity within each lineage (Supplementary Figure 1, Figure 2b) (33). There was good concordance between the diversity represented in the complete data set (Figure 2b) and the derived benchmarking sets (Supplementary Figure 1).

### QuantTB outperforms other tools using simulated data

Using a reference database, QuantTB employs an iterative algorithm to identify which reference strains are present within a sample. Briefly, in every iteration the algorithm selects the most probable genome from the reference database to be present in the sample, based on the number and coverage of SNPs in common between the sample and that particular genome (see Methods). The algorithm outputs the strain(s) predicted to be present within the sample, along with their predicted abundance(s).

We assessed QuantTB’s ability to accurately identify strains across a spectrum of diversity in the reference database using five databases that varied both in size and in the genetic distance between representative genomes (see Methods, Table 1). For testing using these reference databases, we constructed sets of samples containing mixtures of four separate TB strains (see Methods).

We compared QuantTB with two other strain level classification algorithms: Sigma (15) and StrainSeeker (16). However, as Sigma and StrainSeeker are more computationally expensive than QuantTB, we were not able to use our larger databases of mixtures of four strains (>200 strains) with tools other than QuantTB. With larger databases, StrainSeeker could not finish generating its own internal database after seven days, and Sigma took more than 3 days to analyze a single sample, making it impractical or infeasible to benchmark either Sigma or StrainSeeker for larger databases. In contrast, QuantTB scaled well with database size: database construction was complete in less than two hours, and a sample took less than 20 minutes on average to process using the same computer hardware. The ability to take advantage of a large reference database is a substantial advantage for QuantTB over StrainSeeker and Sigma, since the number of publicly available TB sequences in NCBI that could be included in the database is increasing rapidly, and an even larger database. could allow for even finer resolution strain detection.

In order to be able to compare the performance of QuantTB with StrainSeeker and Sigma, we generated a smaller, lower-resolution database of 200 strains (d10small) to assess synthetic mixes of four strains at equal abundance, with coverage levels of either 1× or 10× (See Methods; Table 1). While StrainSeeker performed on par with QuantTB (Figure 3A), both achieving near perfect F1 scores at both coverage levels, Sigma did not perform as well. Sigma identified the correct strains in almost all cases; however, this was accompanied with greatly reduced precision (Supplementary Table 4), i.e. including many false positives and decreasing its overall F1 score (Figure 3A).

**Figure 3).**
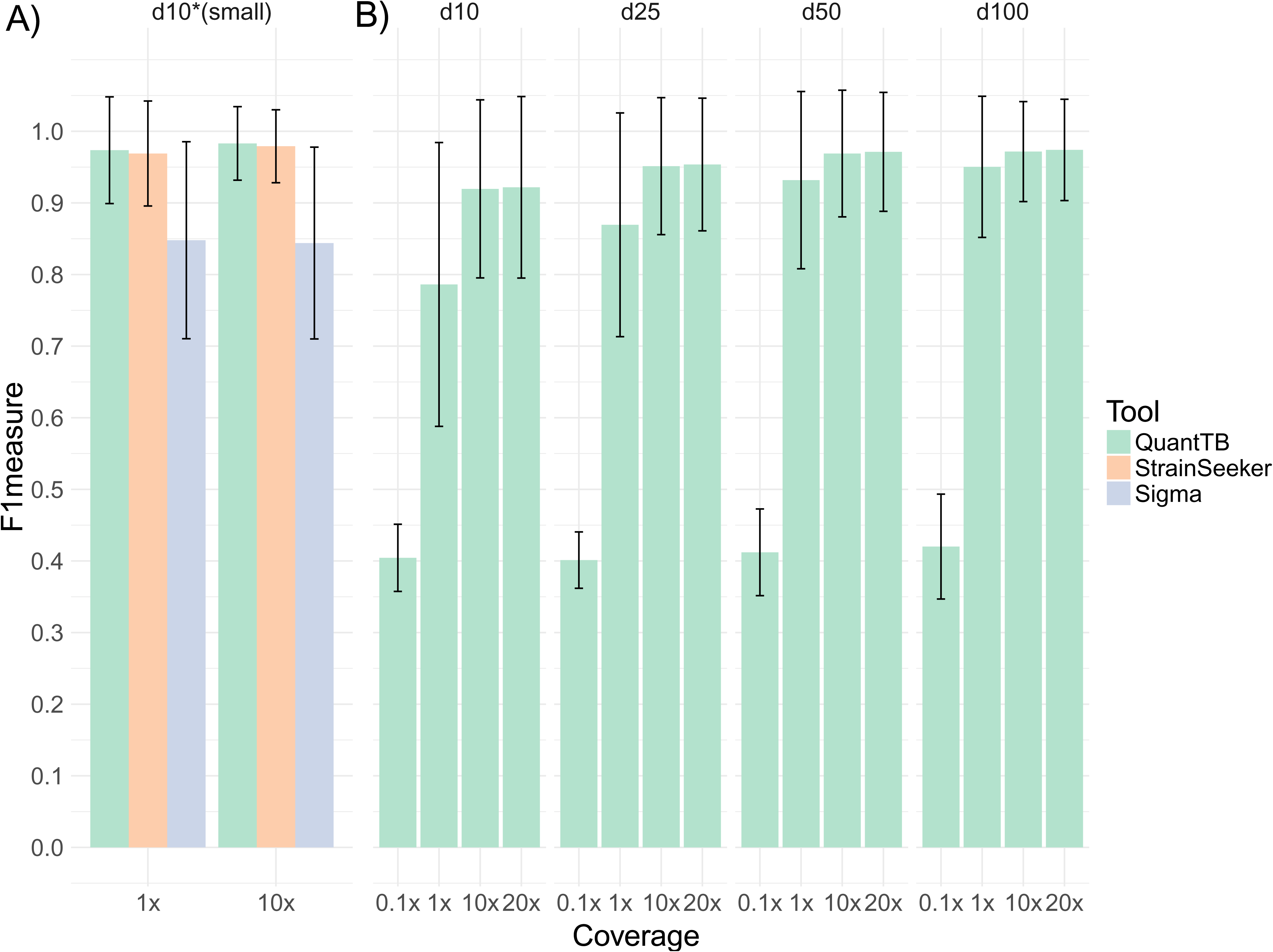
Benchmarking results of synthetically mixed read sets of three different strain identification tools, QuantTB, StrainSeeker and Sigma. A) Results from a smaller database (d10small, n = 200) are shown for all tools for coverage levels of 1× and 10×, B) results from four larger databases (see Table 1) are shown only for QuantTB, for coverages ranging from 0.1× - 20x.

Classification of synthetic four-sample mixes using the larger reference databases presented a more difficult task; however, QuantTB’s performance remained high (Figure 3B), achieving F1 scores above 0.9 at all coverages above 1x per strain, indicating that QuantTB was almost always able to predict all four strains in the synthetic mixes correctly. Scores for lower sample coverage (0.1× per strain) were reduced (F1 score of 0.4). The decreased SNP counts in these very low-coverage simulations led QuantTB to predict only one of the strains present for most of these samples (Supplementary Table 3) We also observed that samples of 20× coverage per strain performed just as well as samples of 10× coverage per strain, indicating no gain in performance from additional coverage. At 1× coverage per strain, QuantTB still performed adequately, with only a slight performance dip noticeable in the largest database containing 4,933 strains differing by at least 10 SNPs. We observed that the lower performance occurred mostly because QuantTB would predict a genetically similar strain instead of the *correct* strain. Taken together, these results suggest that QuantTB can detect strains present at a minimum of 1× coverage. In addition, the fact that the QuantTB algorithm efficiently scales to larger databases not only shows it can accurately classify genomes regardless of database content, but that it runs sufficiently fast to provide the required quick turnaround time in a clinical setting using a large, clinically representative database.

### QuantTB accurately predicts relative abundances

To assess the ability of QuantTB, StrainSeeker, and Sigma to correctly predict relative strain abundances, we simulated mixed samples of pairs of strains that varied in their relative proportions (Figure 4A). As we were only able to use our smallest reference database because of computational limitations with StrainSeeker and Sigma, we used d10small as the reference database to perform these comparisons. To construct test samples for this comparison, we used 100 synthetic mixtures of two assemblies randomly sampled from the d50 database. This represented a more realistic scenario, where strains in the samples (sourced from the d50 database) were not already present in the database (d10small). Pairs of assemblies were combined in ratios of 1:9 and 3:7 relative abundance, with a total sample coverage of 10x.

**Figure 4:**
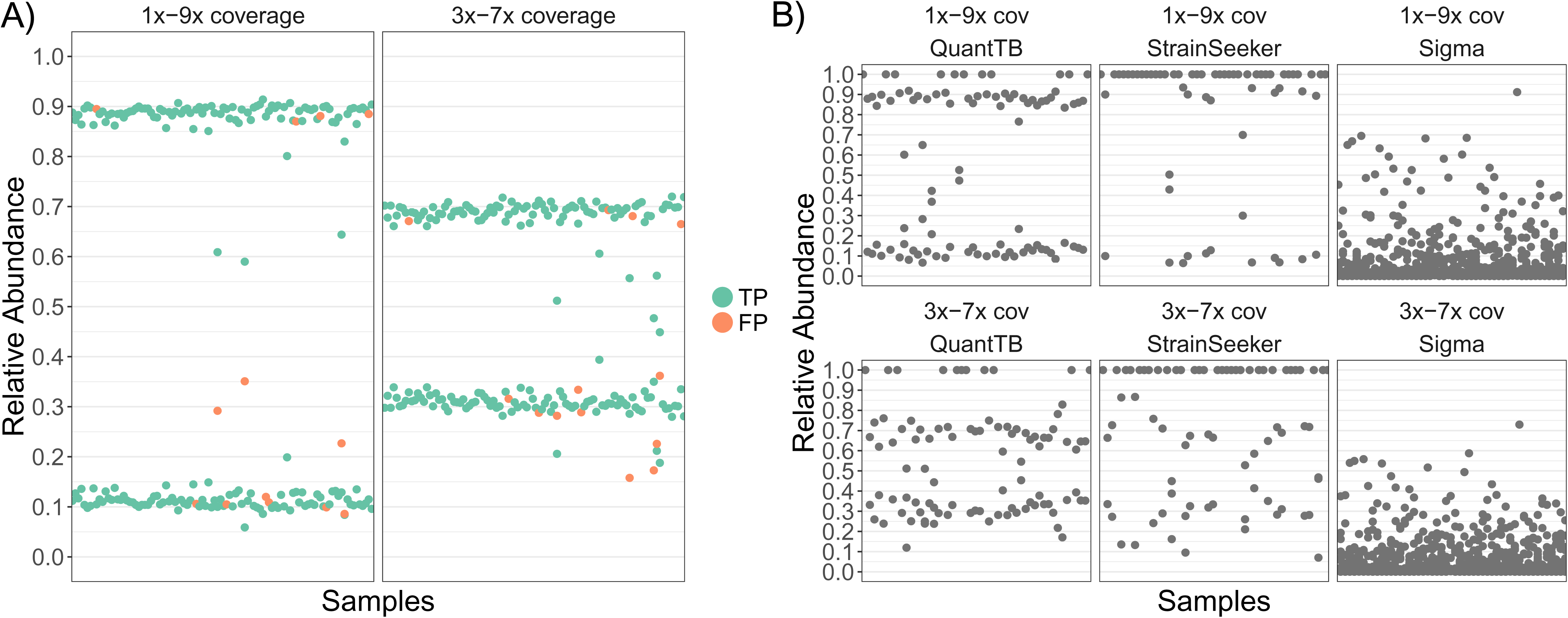
A) Relative abundance predictions across the synthetic sample sets, using randomly selected strains from the d50 and d100 database for QuantTB only. If the strain was correctly predicted for the sample it is colored green (true positive), whereas incorrectly predicted strains are colored red (false positive). The left graph contains samples where two strains are present at 1× and 9× coverage. The right graph contains samples where two strains are present at 3× and 7× coverage. B) Predicted relative abundances across synthetically mixed samples for QuantTB using the d10small database, StrainSeeker and Sigma. Each sample was a pair of strains at either 1×-9× or 3×-7× abundance that were absent from the database. Each point represents a predicted relative abundance for a single strain.

QuantTB was by far the most successful tool at identifying the correct number of strains. QuantTB identified the correct number of strains (two) in the majority of samples (72%). StrainSeeker usually underestimated the number of strains and was only able to identify the correct number of strains in 25% of cases. Sigma failed to predict the correct number of strains in any sample, predicting at least 9 strains for all of the samples (Figure 3B). For samples where QuantTB correctly predicted the strain multiplicity, it also predicted relative abundances close to the expected values, performing best for samples with a 0.1/0.9 strain ratio (Figure 3B, left graphs).

It is not only important to determine whether a tool is able to predict the correct abundances, but also whether it can select the most appropriate genome when the correct strain is absent from the reference database. Therefore, as genomes from the d50 database were used as test samples and tested against genomes in the d10small database, we evaluated the accuracy of strain predictions by assigning a true positive to each strain in a sample if QuantTB predicted the ‘correct’ relative genome in the d10small database (i.e. amongst the top 3 genomes with the highest pairwise SNP distance to the original strain). We found that QuantTB predicts the closest strain to the actual genome with an average precision value of 95%. This more realistic scenario, with previously unseen strains, suggests that QuantTB is able to accurately predict the correct number of strains even in cases where a near-identical strain is not already present in the database. Predictions of Sigma and StrainSeeker for strain multiplicity and relative abundances were insufficiently accurate (Figure 4B and Table 2) to perform this analysis meaningfully.

**Table 2:** Number of samples predicted to contain the specified number of strains, using different methods and databases, for the set of 105 samples from Bryant et al.

| Number of<br>predicted Strains | QuantTB<br>(d10*) | StrainSeeker<br>(d10*) | Sigma<br>(d10*) | QuantTB<br>(d10) | QuantTB<br>(d25) | QuantTB<br>(d50) | QuantTB<br>(d100) |
| --- | --- | --- | --- | --- | --- | --- | --- |
| 1 | 96 | 1 | 1 | 94 | 96 | 95 | 94 |
| 2 | 9 | 0 | 0 | 11 | 9 | 10 | 11 |
| 5 to25 | 0 | 6 | 21 | 0 | 0 | 0 | 0 |
| 26 to 45 | 0 | 90 | 65 | 0 | 0 | 0 | 0 |
| 46 to 60 | 0 | 2 | 8 | 0 | 0 | 0 | 0 |

As only QuantTB could process samples using the larger databases, we further tested its accuracy at identifying correct strain pairs and their differing relative abundances using the d50 and d100 databases. To construct the simulated mixes in our test samples, 100 pairs of assemblies were randomly sampled from both the d50 and d100 databases. As above, pairs of assemblies were combined in ratios of 1:9 and 3:7 relative abundance, with a total sample coverage of 10x. For both databases, QuantTB accurately classified the identity of each strain in the pair (F1 measure of 0.98 and 0.92 for the d100 and d50 databases, respectively, Supplementary Table 4) and accurately determined the relative abundance for each strain in the pair (Figure 4B). The majority of relative abundances predicted were within 0.05 of the correct value (Supplementary Figure 2). Even in the few cases where QuantTB predicted the incorrect strain, QuantTB predicted it to be present in the sample at the correct relative abundance.

### QuantTB differentiates between relapse, reinfection, and mixed infections in real world data

To demonstrate QuantTB’s utility for (clinical) research, we quantified the distribution of *M. tuberculosis* strains within samples from a study investigating the frequency of TB relapses within patients from the REMoxTB clinical trial, a trial which evaluated treatment for TB in previously untreated patients (13). Bryant et al. sequenced 50 pairs of isolates, one taken at an initial time point and the other taken after more than 17 weeks of treatment. Some samples were sequenced more than once (105 total sequencing datasets). Since there are no established methods for detection of mixed infections in *M. tuberculosis* genomic data, the original study used manual inspection of heterozygous SNPs to differentiate between relapse (same infecting strain), reinfection (a different infecting strain) and mixed infections. In the original study, a sample was labeled as mixed if the number of heterozygous loci exceeded a threshold, and as a reinfection if the SNP distance between pairs exceeded a threshold.

Here, we systematically reanalyzed this data using QuantTB and compared our findings from this dataset to those of Sigma and StrainSeeker. As it is impossible to know the identity of the strains present in the real samples in advance, we limited analysis to the multiplicity, or the number of strains identified in each sample. Table 2 shows the multiplicity of infection detected across the dataset of 105 samples for QuantTB, Sigma and StrainSeeker.

QuantTB reported a consistently low (0-2) number of strains, and identified the same seven samples as mixed, irrespective of the database used as a reference, which was in agreement with the expected strain multiplicity based on Bryant et al. In contrast, StrainSeeker and Sigma reported an unrealistically large number of strains (greater than 25 on average).

By applying the results from QuantTB we were able to classify each sample as either part of a relapse, a reinfection or a mixed infection (4 cases). We used results from the d25 database because it performed optimally in our benchmarking tests. If more than one strain was identified by QuantTB, the sample was marked as a mixed infection. If the same strain was identified for both isolates in a pair, the sample pair was marked as a relapse case (35 cases). Finally, if different strains were identified across pairs, the sample pair was marked as a reinfection (3 cases). As Bryant et al. removed some samples due to contamination (in total 9), we also filtered out these samples when comparing our results.

The manual analysis of Bryan et al. designated six samples as mixes. The results from QuantTB match those of Bryant et al for the vast majority of cases (Table 3), classifying the same 3 samples as reinfections, 4 samples as mixed infections, and 33 samples as relapses. QuantTB classified three additional samples as relapses. Samples 42 and 45 were identified as mixed infections in the original study. Upon investigation, it was found that the original study labeled these as mixed infections not based on their original threshold but based off of a ‘manual inspection’, which was not well described. Sample 3 was manually identified as a ‘single isolated positive’, a label given when the second isolate of a pair tested negative for *M. tuberculosis* under culture. Four additional samples were given this label by Bryant et al., who mentioned that these cases were mostly caused by cross contamination. In three culture negative samples labeled ‘single isolate positive’ by Bryant et al., QuantTB identified H37Rv (a laboratory strain). As the coverage for the H37Rv reference strain was high in these three samples, our analysis supports the hypothesis that three culture negative isolates resulting in the sequencing of the H37Rv laboratory strain. The remaining discrepancy, Sample 15, was classified as a reinfection by QuantTB instead of a single isolated positive.

**Table 3:** Comparison of all mixed infections, reinfection and relapses called between QuantTB and Bryant et al. Samples in bold are discordant between the two methods. QuantTB predictions also include the abundance levels of both strains identified within the sample. Samples labeled as Clinically TB negative on follow up were cases in which the second of the isolate pair assigned to the H37Rv strain by QuantTB, and tested negative for TB in the original study.

| Category | QuantTB | Bryant et al. |
| --- | --- | --- |
| Mixed (Lost a strain at later time point) | Sample 2: 81%-18% | Sample 2 |
|  | Sample 8: 53%-47% | Sample 8 |
|  | Sample 23: 60%-40% | Sample 23 |
|  | Sample 50: 75%-25% | Sample 50 |
| Mixed (Gained a strain at later time point) |  | **Sample 42** |
|  |  | **Sample 45** |
| Reinfection | Sample 10 | Sample 10 |
|  | Sample 14 | Sample 14 |
|  | Sample 35 | Sample 35 |
|  | **Sample 15** |  |
| Relapse | 33 matching samples<br>**Sample 42**<br>**Sample 45**<br>**Sample 3** | 33 matching samples |
| Clinically TB negative on follow up | Sample 36 (H37Rv)<br>Sample 37 (H37Rv)<br>Sample 38 (H37Rv) | Sample 36<br>Sample 37<br>Sample 38<br>**Sample 3**<br>**Sample 15** |

To further validate our predictions and clarify discrepancies with the original study, we constructed a phylogenetic tree of all 105 sample isolate pairs based on concatenated SNP sequences (see Methods). This allowed us to visualize the phylogenetic distances between isolates of a sample pair (Figure 5). We observed that most sister leaves in the tree were part of the same sample isolate pair, representing relapse cases. The two samples classified as mixed by the original study but as relapses by QuantTB also appear as sister nodes on the tree (Figure 5, boxes A.1 and A.2). Although this does not rule out a mixed infection, it justifies QuantTB’s relapse classification. In addition, we observed the clustering of isolates which QuantTB identified as most similar to H37Rv (purple nodes in Figure 5, box B), which were classified as ‘single isolated positive’ by the original study. The other samples given this designation by the original study, Sample 3 and Sample 15, did not have an isolate clustered with the H37Rv strain. Instead Sample 3’s isolates were sister nodes on the tree (Figure 5, box C) and the two isolates of Sample 15 were found on opposite ends of the tree (Figure 5, boxes D.1 and D.2), both locations confirm QuantTB’s predictions of relapse and reinfection, respectively.

**Figure 5:**
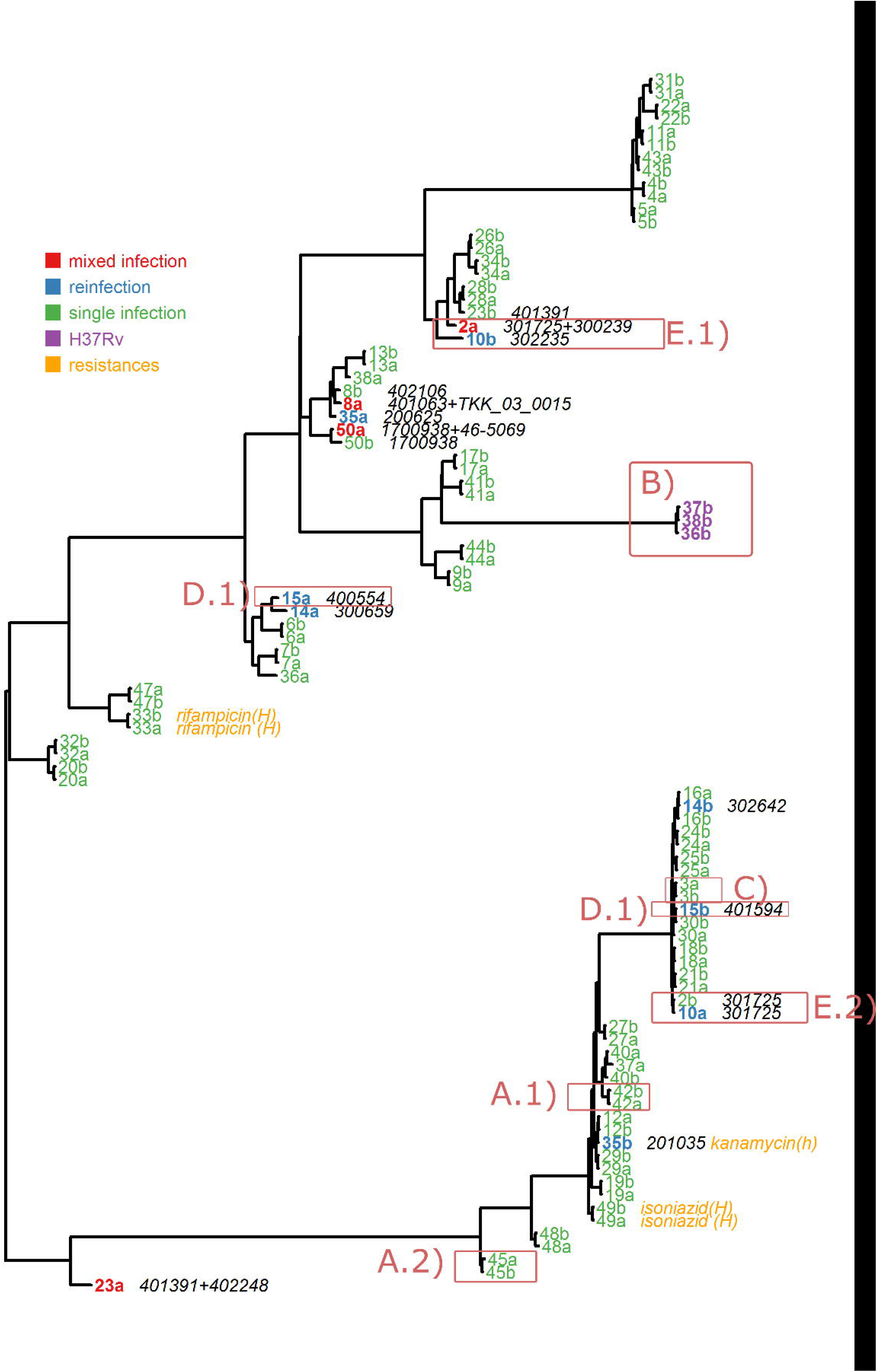
Phylogenetic tree of 47 pairs of isolates from sequencing reads taken from the study of Bryant et al. Tips are labeled with the isolate number and its part of the pair (a or b), and are colored by its isolate classification as predicted by QuantTB. Isolates containing a mixed infection are colored in red. Isolates part of a reinfection pair are colored in blue. Isolates containing the H37Rv strain are colored in purple. Isolates containing antibiotic heterozygous (h) or homozygous (H) resistance mutations are in orange. All single infections isolates are colored in green. To the right of the mixed and reinfection isolates, we show the strains present in the isolate as predicted by QuantTB. Boxes are discussed in the main text.

Finally, we observed two samples whose isolate pairs appeared swapped on the tree: Sample 2 (mixed infection) and Sample 10 (reinfection). Sample 2A has sister nodes with Sample 10B (box E.1), while Sample 10A has sister nodes with Sample 2B on a distant part of the tree (box E.2). Before treatment, Sample 2 (isolate 2A) was mixed with two strains, the minor of which was present within isolate 10A. After treatment, the major strain of Sample 2 was lost, leaving the second pair of Sample 2 (isolate 2B) with only the minor strain, explaining its change of location (next to isolate 10A) on the tree. On the other hand, after treatment, the patient carrying sample 10 was re-infected with a different strain that was similar to the major strain of isolate 2A. Without the annotation of QuantTB it would appear a sample swap might have occurred. But with QuantTB this occurrence can be explained by reviewing the strain identities, because QuantTB outputs which genome has been detected in the sample.

Overall, QuantTB and the manual curation presented in the original study resulted in agreement for 43 of the 47 sample predictions (91%). In the remaining cases, we have presented reasons why QuantTB’s prediction may be at least as accurate as the original manual designations. In addition, QuantTB gives information that was not available from the manual approach of Bryant et al., including detail on multiplicity of infection, and the identity and abundance of each strain, giving a detailed overview of each sample’s genetic makeup.

### QuantTB provides insight into antibiotic resistance

Using QuantTB, we determined the antibiotic resistance genotype for each of the isolates. Antibiotic resistance was indicated if the sample had a SNP in one of the antibiotic resistance causing loci from a previously published curated list (see Methods) (21). Heteroresistance was indicated if the sample had alleles supporting both the resistant and susceptible genotype at a particular locus. Bryant et al. also tested for antibiotic resistance, both phenotypically (with mycobacterial growth indicator tube susceptibility testing) and genotypically (their method was not described). They found no evidence of genotypic or phenotypic antibiotic resistance in any sample. However, we found evidence for genotypic antibiotic resistance in five isolates (Table 4, Figure 5). Two isolates were from the same patient, 33 and 49 (relapse cases) while one was the second isolate in its sample pair, 35b (reinfection case). We found no relation between mixed infections and heteroresistance, nor do we find evidence of the emergence of antibiotic resistance within a relapse case. Isolate 35b exhibited heteroresistance to kanamycin in one locus: 13% of alleles were of the resistance phenotype, and 87% were susceptible. Because this was a reinfection case, it is not possible to determine whether the heteroresistance arose due to within host evolution.

**Table 4:** Isolates exhibiting genotypic antibiotic resistance from the Bryant et al. dataset

| Isolate | Drug | Gene: Amino Acid position, Amino acid change | Type | 599 distribution |
| --- | --- | --- | --- | --- |
| 33a | rifampicin | RpoB: His-445-Ser | homozygous resistant | 600 |
| 33b | rifampicin | RpoB: His-445-Ser | homozygous resistant | 601 |
| 35b | kanamycin | Intergenic: MurA-Ogt | heteroresistant | res: 0.13%<br>sus: 0.87%<br>602 |
| 49a | isoniazid | KatG: Ser-315-Thr | homozygous resistant | 603 |
| 49b | isoniazid | KatG: Ser-315-Thr | homozygous resistant | 604 |

## Discussion

Mixed infections are known to complicate treatment and diagnosis of tuberculosis (8–10); however, the true clinical impact and prevalence of mixed infections is still poorly understood due to the lack of suitable methods to detect and quantify individual strains of *M. tuberculosis*. WGS studies investigating *M. tuberculosis* typically identify mixed infections based on the amount of heterozygous base calls (6,13,14,34). However, both the definition of a heterozygous locus and the number of heterozygous positions indicative of a mixed infection varies between studies. For example, Bryant et al. defined a position as heterozygous when two alleles were supported by at least 5% of the reads with a minimum read depth per allele of 4, and a sample as mixed if it had more than 80 heterozygous base calls (13). Guerra-Assunção et al. defined a position as heterozygous if it had at least 30× coverage and more than one allele accounted for in at least 30% of the reads, and classified a sample as mixed if more than 140 bases were heterozygous (14). Perez Lago et al. simply called a position heterozygous when the less frequent allele was supported by 5 reads (34). With QuantTB we aimed to provide the first algorithm capable of systematically quantifying the multiplicity and abundance of *M. tuberculosis* strains at high resolution using WGS data that does not require manual definitions or counting of heterozygous positions. Because of QuantTB’s unique algorithm that identifies strains in an iterative process, strains can be detected at low coverages (1×), irrespective of the relative frequencies of alleles. The information provided by QuantTB provides several key improvements over a manual approach of counting heterozygous positions. QuantTB: 1) outputs the specific identity of the strain, making the tracking of specific strains across samples possible; 2) outputs the abundances of every strain identified in the sample, enabling the quick identification of major and minor subpopulations; 3) is capable of detecting more than two strains.

Due to QuantTB’s use of a reference database, tracking the presence or absence of specific strains across a set of longitudinal or outbreak samples is also possible. Within a sample, QuantTB can identify the closest strain(s) present from a reference database, even using a large database containing many highly similar genomes (differing by as little as 25 SNPs), allowing us to pinpoint specific strains to within 25 SNPs. This ability to pinpoint (mixes of) specific strains can aid in accurately identifying reinfection cases vs relapses, giving more useful results compared to the manual approach of the Bryant case-study with which our findings are consistent.

Using a systematic approach such as QuantTB aids in identifying cryptic transmission events, such as for samples with dissimilar major strains but matching minor strains. This may have occurred in two of the samples we surveyed in the data of Bryant et al. (samples 2 and 10). The ability to pinpoint strain mixtures can also aid in tracking progression of microevolution between sample isolates, including the evolution of resistance.

Using simulated data, we showed that QuantTB can accurately classify *M. tuberculosis* strains across a variety of database sizes. QuantTB is highly scalable, and can efficiently classify samples with databases as large as 4,000 strains in minutes, a necessary functionality as more and more TB assemblies are resolved. Other published tools made for classifying single strains in samples, StrainSeeker and Sigma, were not capable of working with large databases, limiting their applicability as a diagnostic tool for *M. tuberculosis*. On tests using a smaller database - an easier and low resolution experiment - QuantTB identified the strain composition of synthetic sets with comparable accuracy as StrainSeeker, while Sigma’s results included numerous false positives. On tests where the mixed samples contained strains absent from the database, QuantTB outperformed the other tools by accurately outputting the correct multiplicity in 72% of cases, in comparison to 25% for StrainSeeker and 0% for Sigma. Both Sigma and StrainSeeker consistently outputted aberrantly high number of strains. In addition, QuantTB predicted the closest related genome in the database for these strains in 94% of the samples.

The detection of high quality SNPs in a sample is an essential part of QuantTB’s algorithm. In order to ensure erroneous SNPs are not considered, QuantTB disregards SNPs present at less than 5% abundance relative to that of the previously identified strain. Therefore, QuantTB can only detect mixed infections in which the minor strain represents at least 5% of the allelic variation. However, QuantTB is still able to pinpoint low-abundance strains with greater sensitivity than previous approaches based on the counting of heterozygous positions, due to its ability to identify strain down to coverages as low as 1x.

An advantage of approaches based purely on heterozygous locations is that they do not depend on a reference database. QuantTB’s ability to accurately detect mixed infections is closely integrated with the distribution of genomes used to construct the database. Though we have tested QuantTB’s performance on samples containing strains absent from the database, we have not extensively tested how the absence of a large proportion of a strain’s lineage would affect its classification. QuantTB’s ability to detect a strain not in the database depends on how distant it is from its nearest relative in the database. If the strain is sufficiently distant, it is likely that the strain would not be detected, underestimating sample diversity. The effects of QuantTB’s database reliance is mitigated by ensuring the database covers as much diversity as possible. We found the currently available data is skewed to favor genomes of lineage 4 and lineage 2, with lineage 7 and 5 representing only 0.2% of the downloaded assemblies. Therefore, further sequencing of these underrepresented lineages would aid QuantTB in proper classification of novel strains.

QuantTB determines antibiotic resistance phenotypes by querying the sample against a manually curated list of SNPs that were shown to cause antibiotic resistances in previous studies. Bryant et al. did not find clinical evidence for antibiotic resistance amongst the samples. Using the curated list we found antibiotic resistance in five samples, one being a case of heteroresistance in the second isolate of its sample pair. We did not observe any relationship between antibiotic resistance and mixed infections in the clinical isolates. The observed resistance mutations are well-known causal mutations for their respective resistances and WGS has been shown to outperform phenotypic susceptibility tests for predicting resistance (35). Because Bryant et al. did not mention which method of genotypic testing they employed, it is not possible to understand why they were unable to detect genotypic resistances in the isolates. Particularly the *katG* mutation predicted from genotypic data in samples 49a and 49b is widely known and confirmed to confer resistance to isoniazid. The ability to accurately determine antibiotic resistance from sequencing data is still an active research topic for TB (36, 37). As antibiotic resistance is one of the biggest threats to world-wide TB eradication, the proper detection of possible resistance in samples is crucial.

## Conclusion

We introduce QuantTB, a new classification method that leverages the high-resolution capability of whole genome sequencing for the detection of mixed *M. tuberculosis* infections. In contrast to existing tools such as Sigma and StrainSeeker, QuantTB is scalable and able to leverage a high-resolution reference database representing the scope of diversity within TB. Even when using a smaller database that allows comparisons between these tools, QuantTB shows substantially better performance on both synthetic and clinical datasets. This tool can be used to rapidly and accurately identify specific *M. tuberculosis* strains in clinical samples, track transmission of TB strains across longitudinal samples and outbreaks, and differentiate between relapse and reinfection cases. The ability to disentangle mixed infections in an accurate and scalable manner will help control TB and help limit the spread of antibiotic resistance.

## Declarations

### Ethics approval and consent to participate

Not applicable

### Consent for publication

Not applicable

### Availability of data and material

The FASTA files used in this study can be downloaded from NCBI using the accession numbers listed in Supplementary Table 1.

The raw sequence data analyzed in this study can be downloaded from the Sequencing Read Archive (BioProject Accession: PRJEB2777).

QuantTB can be downloaded and installed from github: https://github.com/AbeelLab/quanttb/

## Acknowledgements

### Authors’ contributions

AK, TA, AE, BW conceived the project. CA and TA designed and implemented the analyses and wrote the manuscript. AM, TS, TA, AK, AE provided comments on the manuscript. All authors read and approved the final manuscript.

